# Overexpression of Orange (OR) and OR mutant protein in *Chlamydomonas reinhardtii* enhances carotenoid and ABA accumulation and increases resistance to abiotic stress

**DOI:** 10.1101/2020.05.10.087080

**Authors:** Mohammad Yazdani, Michelle G. Croen, Tara L. Fish, Theodore W. Thannhauser, Beth A. Ahner

## Abstract

The carotenoid content of plants can be increased by overexpression of the regulatory protein ORANGE (OR) or a mutant variant known as the ‘golden SNP’. In the present study, transgenic lines of the microalgae *Chlamydomonas reinhardtii* were generated to overexpress either wild type *CrOR* (*CrOR^WT^*) or a mutated *CrOR* (*CrOR^His^*) containing a single histidine substitution for a conserved arginine. Overexpression of both *CrOR^WT^* and *CrOR^His^* dramatically enhanced the accumulation of several different carotenoids, including β-cartotene, α-carotene, lutein and violaxanthin, in *C. reinhardtii* and, in contrast to higher plants, upregulated the transcript abundance of several relevant carotenoid biosynthetic genes. In addition, microscopic analysis revealed that the *OR* transgenic cells were larger than control cells and exhibited larger chloroplasts with a disrupted morphology. Moreover, both *CrOR^WT^* and *CrOR^His^* cell lines showed increased tolerance to salt and paraquat stress. The levels of endogenous phytohormone abscisic acid (ABA) were also increased in *CrOR^WT^* and *CrOR^His^* lines, not only in normal growth conditions but also in growth medium supplemented with paraquat. Together these results offer new insights regarding the role of the OR protein in regulating carotenoid biosynthesis and accumulation in microalgae, and establish a new functional role for *OR* to modulate oxidative stress tolerance mediated by ABA.

## 1. Introduction

Carotenoids are valuable secondary metabolites, vital to many life forms, but only synthesized in plants, fungi, algae, and bacteria. In humans, carotenoids are an essential part of the diet, providing pro-vitamin A and reducing the onset of chronic diseases such as cardiovascular diseases, cancers, and other age-related diseases (Fraser and Bramley, 2004; Fiedor and Burda, 2014). In aquaculture and poultry industries, carotenoids such as β-carotene, astaxanthin, canthaxanthin, lycopene, lutein, and zeaxanthin are used as a feed supplementation to enhance the color of egg yolk, chicken, and fish meat (Vílchez et al., 2011; Sathasivam and Ki, 2018). Due to all aforementioned features of carotenoids, the demand and market for these natural products are growing remarkably (Sathasivam and Ki, 2018). They are largely sourced from plant and algal (Takaichi, 2011; Sun et al., 2018).

In plants and algae, carotenoids function as accessory pigments in light-harvesting complexes during photosynthesis. They protect photosynthetic machinery from photooxidative damage by scavenging reactive oxygen species (ROS) (Niyogi, 1999; Vílchez et al., 2011; Domonkos et al., 2013). They are also precursors for the production of volatiles, phytohormones, and signaling molecules (Umehara et al., 2008; Alder et al., 2012; Avendaño-Vázquez et al., 2014).

Carotenoids are synthesized in plastids via the methylerythritol 4-phosphate (MEP) pathway, which provides the 5C building blocks isopentenyl diphosphate (IPP) and its isomer dimethylallyldiphosphate (DMAPP) (Eisenreich et al., 2004; Nisar et al., 2015). As shown in Figure 1, both IPP and DMAPP can be converted to geranylgeranyl diphosphate (GGPP) and two molecules of GGPP are condensed to form phytoene. This is the first committed step for carotenoid biosynthesis and is mediated by the rate-limiting enzyme phytoene synthase (PSY) which directs the metabolic flux towards carotenogenesis (Lu et al., 2006; Tzuri et al., 2015). Phytoene is then converted into lycopene via subsequent steps of desaturation and isomerization. Lycopene can then be cyclized by cyclases to produce α- and β-carotene. Further hydroxylation and epoxidation result in the production of xanthophylls such as violoxanthin and neoxanhtin which can be converted to ABA by 9-*cis*-epoxycarotenoid dioxygenase (NCED) (Ruiz-Sola and Rodríguez-Concepción, 2012; Nisar et al., 2015; Yuan et al., 2015b).

**Figure 1:**
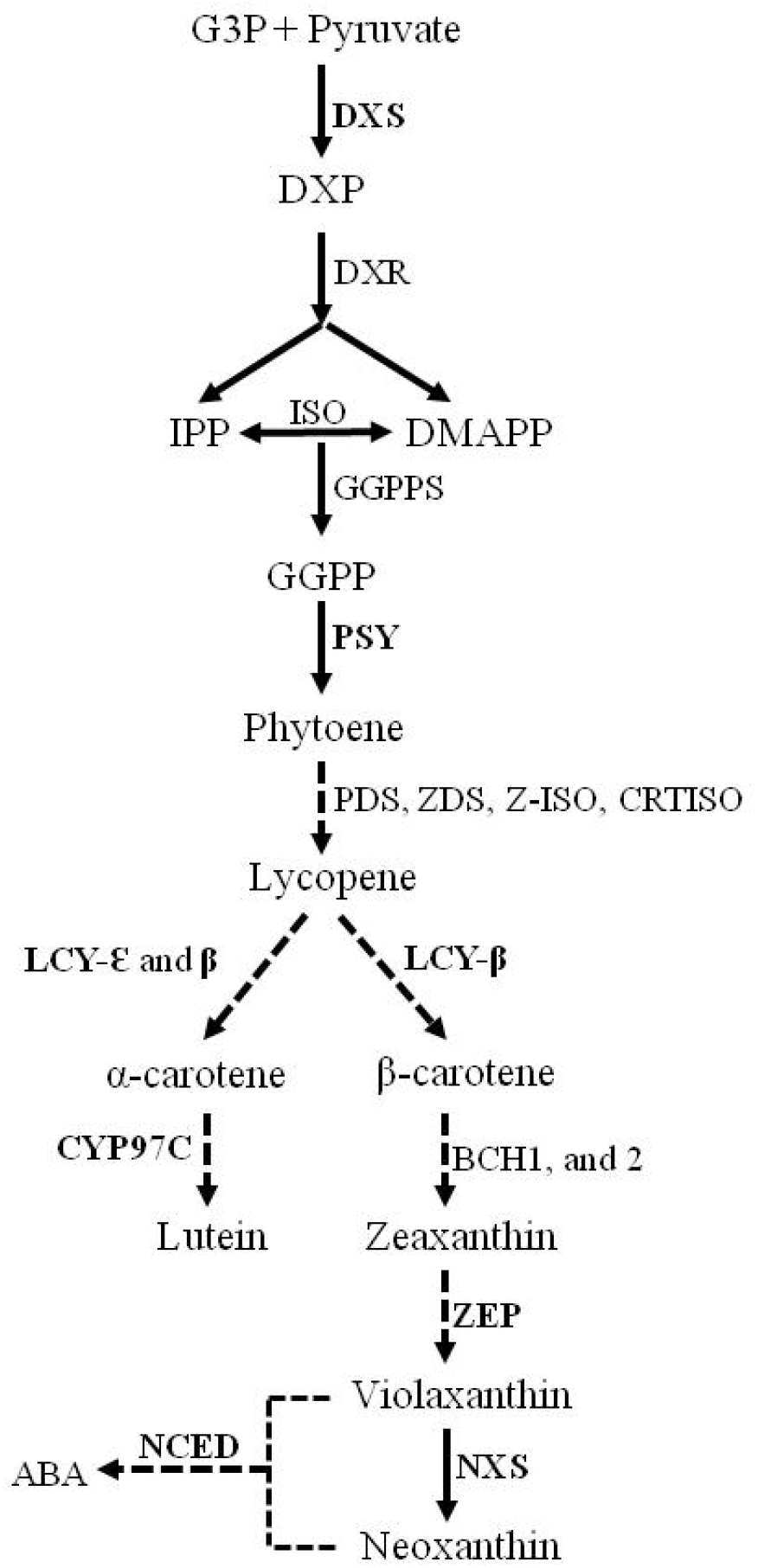
Carotenoid biosynthesis pathway in plants. Dashed arrows display multiple steps. G3P, glyceraldehyde-3-phosphate; DXP, 1-deoxy-D-xylulose-5-phosphate; DXS, 1-deoxy-D-xylulose-5-phosphatesynthase; ISO, isomerase; DXR, 1-deoxy-D-xylulose 5-phosphatereductoisomerase; IPP, isopentenyl diphosphate; DMAPP, dimethylallyl diphosphate, GGPP, geranylgeranyldiphosphate; GGPPS, geranylgeranyl diphos-phate synthase; PSY, phytoene synthase; PDS, phytoene desaturase; ZDS,z-carotene desaturase; Z-ISO, ζ-carotene isomerase; CRTISO, carotenoid isomerase; LCY-□, lycopene ε-cyclase; LCY-β, lycopene β-cyclase; CYP97C, cytochrome P450-type monooxygenase 97C; BCH1 and 2, β-carotenoid hydroxylase 1and 2; ZEP, zeaxanthin epoxidase; NXS, neoxanthin synthase; NCED, 9-*cis*-epoxycarotenoid dioxygenase; ABA, abscisic acid. The genes corresponding to the enzymes shown in bold were used for q-PCR.

Strategies to elevate the carotenoid content in plants have focused on altering the gene expression of key carotenogenic enzymes. For instance, down-regulation of lycopene-*ε*-cyclase (*LCY-ε*) in potato (Diretto et al., 2006) and tobacco (Shi et al., 2014) increased total carotenoids by 2 to 2.5-fold. Simultaneous down-regulation of *LCY-ε* and *β*-carotene hydroxylase (*CHY-β*) increased total carotenoids up to 2.7-fold in sweet potato plants and 18-fold in calli (Kim et al., 2012; Ke et al., 2019). Overexpression of bacterial *PSY* in tomato fruits increased total carotenoids by 4-fold (Fraser et al., 2002). A CRISPR/Cas9-based approach was also successfully used in banana fruit to edit the *LCY-ε* gene for targeted increase of *β*-carotene (Kaur et al., 2020).

Plants have been shown to utilize the ORANGE (OR) protein to regulate carotenoid accumulation and chromoplast biogenesis (Lu et al., 2006; Tzuri et al., 2015). Highly conserved in plant species, the *OR* encodes a DnaJ cysteine-rich zinc finger domain-containing protein that is targeted to plastids (Lu et al., 2006). Native OR is known to physically interact with PSY in *Arabidopsis*, melon, and sweet potato, and is required to sustain PSY protein levels for carotenoid biosynthesis (Zhou et al., 2015; Park et al., 2016; Chayut et al., 2017). Overexpression of wild type *OR* resulted in high levels of carotenoids in various organs including sweetpotato storage roots, *Arabidopsis* leaves, rice grains, and corn endosperm (Park et al., 2015; Yuan et al., 2015a; Bai et al., 2016; Berman et al., 2017). Functional analysis of the *OR* genes in various orange-flesh melon fruits identified a single-nucleotide polymorphism known as ‘golden SNP’, which converts a highly-conserved arginine to histidine and is responsible for elevated β-carotene accumulation (Tzuri et al., 2015). Overexpression of the altered *Arabidopsis OR* (*AtOR^His^*), result in a greater increase in carotenoid levels relative to controls compared to overexpression of the wild type *OR* (*AtOR^WT^*) in *Arabidopsis* calli and in tomato fruit (Yuan et al., 2015a; Yazdani et al., 2019).

Because microalgae such as *Dunaliella salina* and *Haematococcus pluvialis* naturally accumulate high levels of carotenoids (Cezare-Gomes et al., 2019) and other microbes such as yeast can be genetically modified to produce carotenoids (Cataldo et al., 2020), microbial production is viewed as an efficient way to produce these valuable compounds. Carotenoid biosynthesis has therefore also been studied in *C. reinhardtii*, a genetically tractable well-characterized model algae species (Lohr et al., 2005; Takaichi, 2011). The basic carotenoid biosynthetic genes are highly conserved between plants and *C. reinhardtii*, although some algae species are capable of synthesizing unusual carotenoids through specialized pathways (Takaichi, 2011; Varela et al., 2015). As with higher plants, targeted alteration of specific carotenoids in algae has mainly involved altering gene expression of enzymes involved in carotenoid biosynthesis and/or ectopic expression of a carotenogenic gene from another species. For instance, overexpression of phytoene desaturase in *H. pluvialis* increased total carotenoid levels by about 30% (Steinbrenner and Sandmann, 2006). Overexpression of β-carotene ketolase gene from *H. pluvialis* in *C. reinhardtii* resulted in the production of new carotenoid keto-lutein (León et al., 2007), and overexpression of the *PSY* gene from *D. salina* in *C. reinhardtii* led to an increase of 125% in lutein and 260% in *β*-carotene content (Couso et al., 2011).

Originally annotated as *CPL6*, the *OR* gene is also present in *C. reinhardtii* (Merchant et al, 2007; Sun et al., 2010). The wild type *C. reinhardtii* OR (*CrOR^WT^*) protein sequence consists of 302 amino acids and shows 56.7% identity to the AtOR protein sequence (KEGG: Kyoto Encyclopedia of Genes and Genomes). In a previous study, *CrOR^WT^* overexpression in *C. reinhardtii* increased lutein and β-carotene levels by 1.9- and 1.7-fold per cell, respectively, using dual promoters of heat-shock protein 70A and ribulose bisphosphate carboxylase small chain 2 (Morikawa et al., 2018). However, *CrOR* transcript and protein levels were not reported in this study nor were the levels of other carotenoids. In the present study, we used a strong light-inducible promoter to overexpress *CrOR^WT^* and the mutated *CrOR* gene *CrOR^His^* in *C. reinhardtii* and analyzed the physiology, biochemical composition, and expression of some genes in the resulting mutant strains. Expression of the *OR* transgenes significantly boosted the total carotenoid levels in *C. reinhardtii* cells. In addition, *CrOR^WT^* and *CrOR^His^* transformants were better able to tolerate abiotic stress caused by salt and paraquat, possibly due to increased ABA levels. Moreover, overexpression of the *OR* genes noticeably altered *C. reinhardtii* cell size and chloroplast shape. These results demonstrate that changes in the *OR* expression levels and/or in the OR amino acid sequence are important factors that may be used to increase pigment levels in algae for commercial purposes and to confer resistance against various abiotic stress conditions.

## 2. Material and methods

### 2.1. Algae strain and growth conditions

*C. reinhardtii* cell-wall mutant CR4349 strain and the derived transgenic lines were maintained in TAP medium (20 mM Tris, 17 mM Acetate, 0.68 mM K_2_HPO_4_, 7.26 mM KH_2_PO_4_, and 7.5 mM NH4Cl) (Togasaki et al., 1987) with the addition of trace metals as described previously (Hunter, 1946). Experiments were carried out in 50-400 mL batch cultures of TAP medium incubated at 25 °C with constant shaking (130 rpm), open to the atmosphere and under continuous fluorescent white light (300 μmol m^-2^ s^-1^) unless otherwise stated. The algal cell growth was monitored by measuring the auto-fluorescence of chlorophyll using a 10-AU Fluorometer (Turner Designs, San Jose, CA), equipped with a Daylight White Lamp F4T5D (EX: 340–500 nm, Em: >□665 nm) (Richter et al., 2018).

### 2.2. Site-directed mutagenesis, plasmid construction, and algae transformation

*C. reinhardtii* OR protein (*CPL6*, Phytozome ID number Cre06.g279500) and its conserved motif containing arginine-106 (Arg106) was identified by alignment with *Arabidopsis thaliana* OR (At5g61670) and melon (*Cucumis melo*) CmOR (GenBank accession numbers KM505047 and KM505046 for green and orange flesh melon, respectively) (Tzuri et al., 2015). The *CrOR* open reading frame was first PCR amplified from *C. reinhardtii* cDNA along with the addition of a FLAG tag to its C-terminus, and then cloned into SacI and XmaI sites of PUC19 plasmid cloning vector. Site-directed mutagenesis was performed to change Arg106 to a histidine (His) using Q5^®^ Site-Directed Mutagenesis Kit (New England Biolabs Inc.) according to the manufacturer’s instructions. The *CrOR^WT^* and *CrOR^His^* amended with a FLAG tag added to their C-termini, were then sub-cloned into the SalI site of the pCAMBIA1300 vector. The *PsaD* promoter and terminator was amplified from *C. reinhardtii* as previously described (Fischer and Rochaix, 2001; Kumar et al., 2013), and cloned into the SacI/KpnI and PstI/HindII sites of the transformation vector, respectively. All primers are shown in Supplementary Table 1. *C. reinhardtii* (cell-wall mutant CR4349 strain) transformation was carried out using the glass bead method as previously described (Kindle, 1990) and the positive single colony of transformants grown on TAP plates supplemented with 10 mg.L^-1^ hygromycin B were used for the experiments.

### 2.3. Protein extraction and immunoblotting

All *C. reinhardtii* cultures were grown in TAP medium and harvested by centrifugation (7,500 rpm, 15 min) at exponential growth phase. Pellets were then resuspended in 1 mL of 10 mM sodium phosphate buffer pH 6.8 with 1 mM PMSF as proteinase inhibitor (Díaz-Troya et al., 2008). Resuspended pellets were subjected to probe sonication with 20/40 s on/off duty cycle on ice. The cell debris was separated from the supernatant by micro-centrifugation at 13,000 rpm for 10 min at 4 C°. The soluble fraction was used to measure total protein using the Bradford method (BioRad, Hercules, CA).

For immunoblotting, the total soluble proteins were separated by 12% SDS-PAGE gels and blotted onto nitrocellulose membrane. The membrane was then blocked by 5% (w/v) nonfat milk in TBST (0.1 M Tris, pH 7.4, 0.15 M NaCl, 0.1% Tween 20) and incubated with monoclonal FLAG tag antibody (ThermoFisher Scientific). Commercial goat anti-rabbit IgG antibody (Invitrogen) was used as the secondary antibody. Signals were detected by ECL Western Blotting Substrat (Promega, USA).

### 2.4. Carotenoid extraction and analysis

Carotenoids were extracted from *C. reinhardtii* cell pellets using 80% acetone (v/v) as previously described (León et al., 2005). Samples were run in an injection solvent of ethyl acetate containing 0.05mg /ml DIM (diindolylmethane) as an internal standard. Carotenoids were separated within 12 minutes on a Waters UPC2 (ultra-performance convergence chromatography) system with a Viridis HSS C18 SB column (3.0 × 100 mm, 1.8 μm) at 40 °C and a flow rate of 1 ml/min using gradient elution with a two component mobile phase system of supercritical carbon dioxide (SC-CO2) and methanol (MeOH). The column was equilibrated at 1% MeOH. The analytes were eluted using a non-linear concave gradient to 20% MeOH over 7.5 minutes. The mobile phase composition was held constant at these values for 4.5 minutes. The column was then re-equilibrated to initial conditions over 3 minutes (Supplemental Figure S1 and Supplemental Table S2). The automatic back pressure regulator (ABPR) setting was varied throughout the run as described in Supplemental Table S3. UV/VIS detection of carotenoid peaks was performed on a Waters photodiode array (PDA) detector over a range of 250nm to 700nm. Each of the subject carotenoids was quantified at unique wavelengths optimized for both sensitivity and selectivity (Supplemental Table S4). The β-carotene was used as the calibration standard. The concentrations of all other carotenoids are reported in β-carotene equivalents using the TargetLynx software in MassLynx 4.1 (Waters Milford, MA) (Shinozaki et al., 2018).

### 2.5. Confocal microscopy

The autofluorescence of carotenoids and chlorophyll *a* was observed in cell-wall mutant and *OR* transgenic cells during exponential growth phase using Leica TCS SP5 Laser Scanning Confocal Microscope. A 488 nm argon laser line was used as the excitation source and the detector set at 650-700 nm and 500-550 nm for chlorophyll *a* and carotenoids, respectively (D’Andrea et al., 2014). The Image J software was used to measure the projected cell surface area and estimate the volume of the cells.

### 2.6. RNA extraction and qRT-PCR

After centrifugation of a 50 ml aliquot of the algae cell culture at exponential growth phase, the total RNA from the cell pellet was extracted using RNeasy Plant Mini Kit according to manufacturer’s manual (Qiagen, Carlsbad, CA, USA). 1 μg of total RNA was used to synthesize cDNA using iScript™ cDNA Synthesis Kit (Bio-Rad). Real-time RT-PCR was performed using gene specific primers (Supplementary Table 1) and SYBR Green Supermix (Bio-Rad) in a CFX96 Real-Time PCR system (Bio-Rad) as described (Zhou et al., 2011). The *C. reinhardtii rbcL* gene was used as an internal control and the relative changes in expression levels were analyzed using the 2^−ΔΔCT^ method (Lyi et al., 2007). All experiments were carried out with three biological replicates.

### 2.7. Salt and paraquat stress experiments

For all stress experiments, cultures containing the same cell density were prepared in the tubes containing 20 ml of TAP medium supplemented with various concentrations of NaCl (0.5, 0.1, and 0.2 M) and paraquat (5, 15, and 50 nM). The tubes were then incubated at 25 °C with shaking manually once a day under continuous fluorescent white light (300 μmol m^-2^ s^-1^). The algae cell growth was monitored daily by measuring the auto-fluorescence of chlorophyll *a* using a 10-AU Fluorometer (Turner Designs, San Jose, CA), equipped with a Daylight White Lamp F4T5D (EX: 340–500 nm, EM: >□665 nm) (Richter et al., 2018).

### 2.8. ABA extraction and quantification

To measure the ABA concentration, the cell-wall mutant and transgenic lines were grown in flasks containing 400 ml TAP medium supplemented with or without 15 nM paraquat. The flasks were incubated at 25 °C with constant shaking (130 rpm) under continuous fluorescent white light (300 μmol m^-2^ s^-1^). The cells were harvested via centrifugation during exponential growth phase, washed twice with deionized water, and freeze dried using a FreeZone 2.5 Liter lyophilizer (LABCONCO^®^, USA). ABA was measured in extracts prepared using 300-500 mg of the lyophilized cells and the ABA ELISA Kit as described by manufacturer’s instructions (MyBioSource).

### 2.9. Statistical analysis

Data analysis was performed using Student’s *t* test (Suzuki et al., 2008) to show the significance of differences between data sets.

## 3. Results

### 3.1. Overexpression of CrOR^WT^ and CrOR^His^ genes in C. reinhardtii

The coding sequence of the wild type *OR* gene (*CrOR^WT^*) in the *C. reinhardtii* was identified using the *A. thaliana OR* as a reference gene and the conserved motif containing arginine (R106) was found by aligning the *C. reinhardtii* OR protein sequence against various plant species (Figure 2A) including melon protein CmOR. Site-directed mutagenesis was used to change R106 to a histidine creating *CrOR^His^*. Overexpression of *CrOR^WT^* and *CrOR^His^* genes in *C. reinhardtii* was driven by the strong endogenous promoter from the *PsaD* gene (Figure 2B), which is both constitutive and light-inducible and controls the expression of an abundant chloroplast protein in photosystem I (Fischer and Rochaix, 2001; Kumar et al., 2013). The expression of *OR* gene and OR protein levels were confirmed by q-PCR and immunoblot analysis, respectively. The results showed similarly high expression levels of both OR transcript and protein in all tested *CrOR^WT^* and *CrOR^His^* lines (Figure 2 C and D).

**Figure 2:**
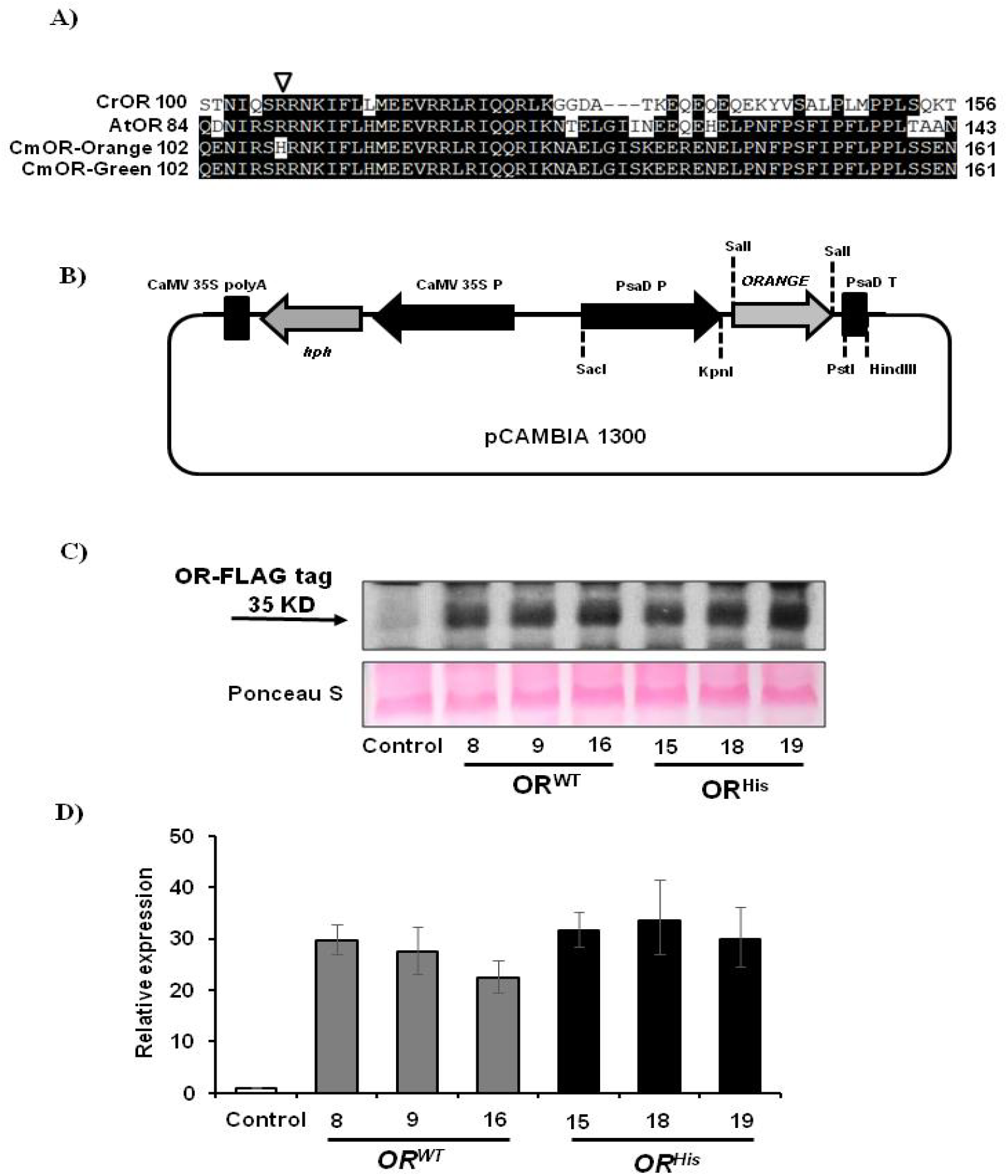
Characterization of *CrOR* transgenic lines. A, Alignment of partial OR amino acid sequences from *C. reinhardtii* (CrOR), *Arabidopsis* (AtOR), melon (CmOR) with green and orange flesh. The triangle shows the arginine (R) to histidine (H) mutation in orange flesh melon. B, Simplified diagram depicting *ORANGE* gene cassette in pCAMBIA 1300 vector (P: promoter, T: terminator). C, Immuno-blot analysis of OR protein level in *C. reinhardtii*. D, qRT-PCR analysis of the *OR* gene expression in *C. reinhardtii* transgenic cells.

### 3.2. Overexpression of the CrOR^WT^ and CrOR^His^ promotes carotenoid accumulation in C. reinhardtii

The levels of various carotenoids were quantified by UPC^2^ in control and transgenic *C. reinhardtii* and reported on a per cell basis. The total carotenoid levels in all three *CrOR^His^* overexpressing lines were significantly higher than those in both control and *CrOR^WT^* overexpressing lines (Figure 3 and Figure 5A). The *CrOR^His^* line 15 displayed the largest difference in total carotenoids compared to the other lines. The total carotenoid content in this line was 1.3- and 3-fold higher than the high-carotenoid *CrOR^WT^* line 16 and control, respectively (Figure 3). These results suggest that overexpression of both *CrOR^WT^* and *CrOR^His^* promote carotenoid biosynthesis and accumulation in *C. reinhardtii* but *CrOR^His^* is significantly more effective.

**Figure 3:**
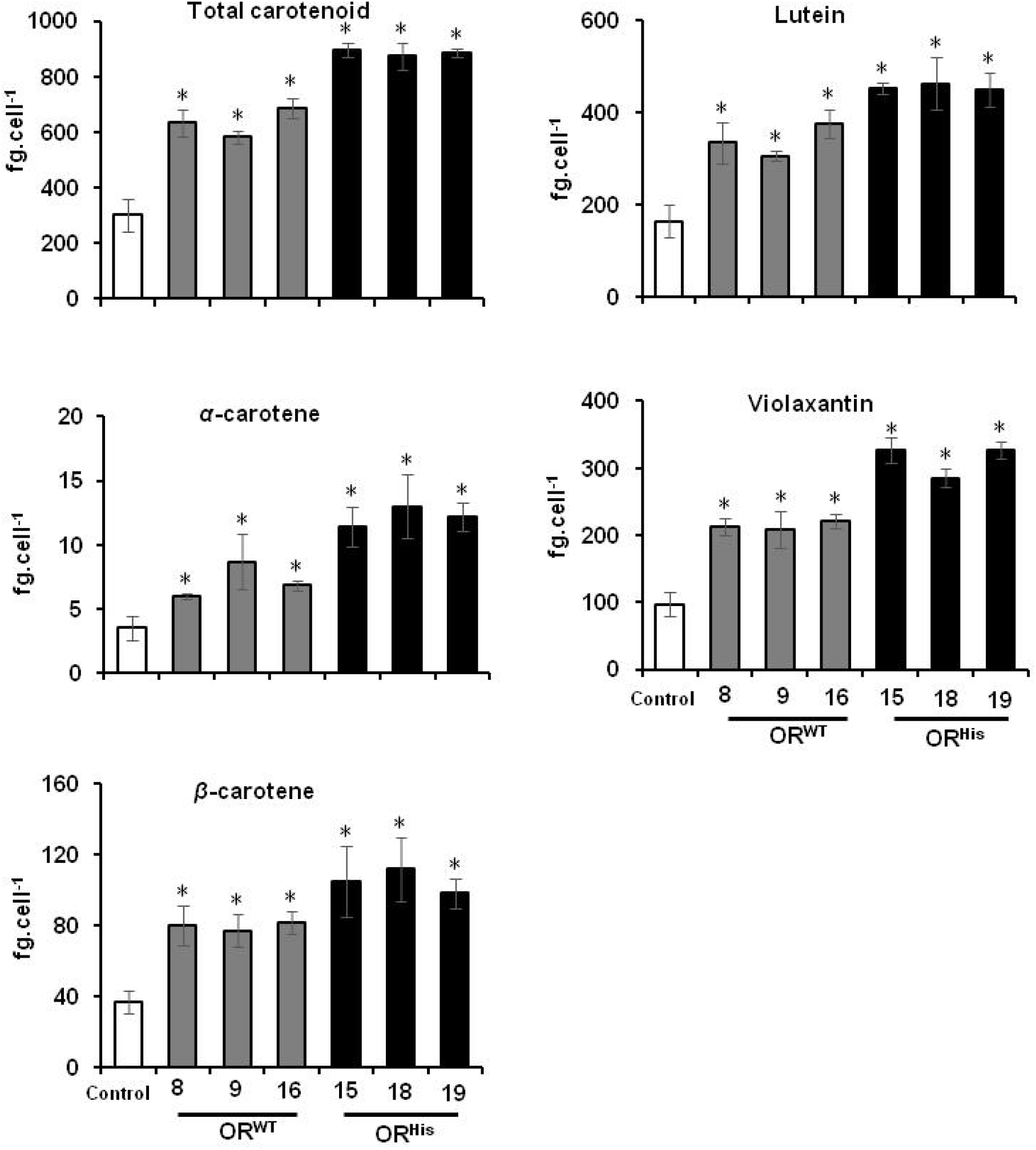
Total carotenoid and carotenoid profile, and levels in *OR* transgenic cells. Data are the mean of three biological replicates ± SD. * indicates significant difference at *P*<0.05.

**Figure 4:**
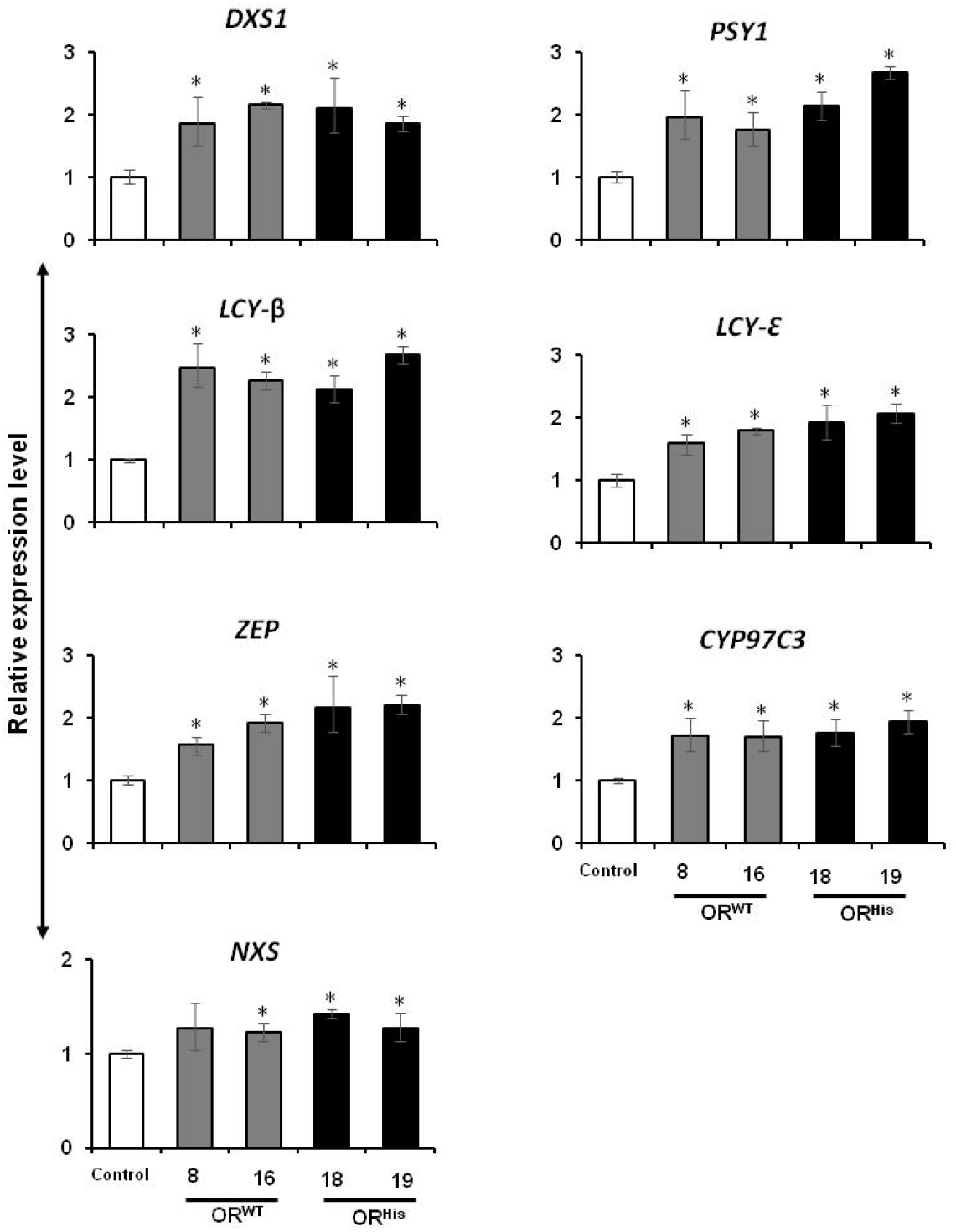
Transcript levels of carotenogenic genes in *CrOR* transgenic lines. Data are the mean of three biological replicates ± SE. * indicates significant difference at *P*<0.05.

**Figure 5:**
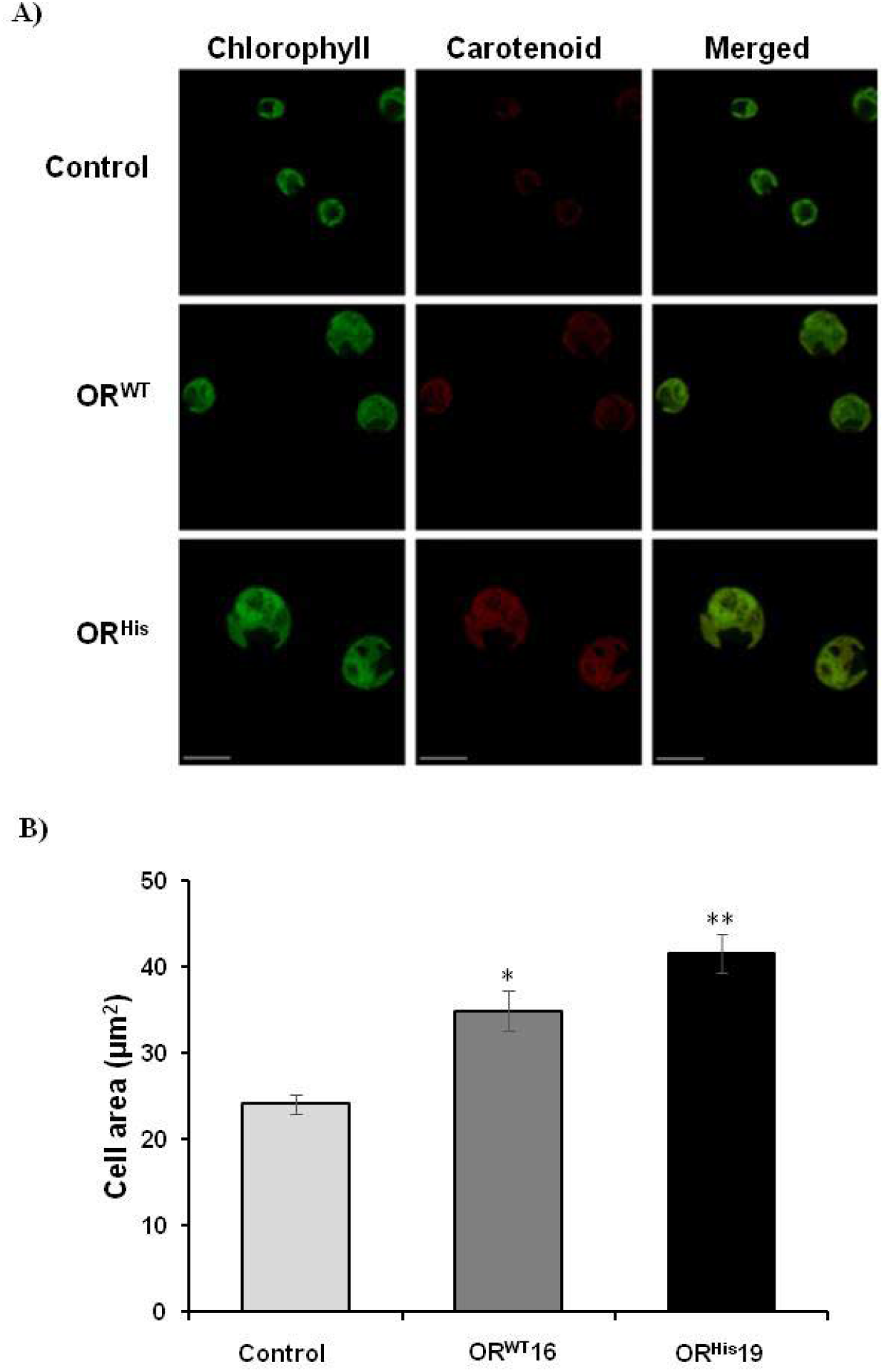
*CrOR* overexpression impact on transgenic cell lines. A, autofluorescence of carotenoids and chlorophylls (set as red and green signal, respectively) in algae transformants using confocal microscope. The size of the bars is 10 μm. B, projected cell surface area in *OR* transgenic lines. Data are the mean of three biological replicates (20-30 cells per replicate) ± SD. * indicates significant difference at *P*<0.05.

Among the various forms of carotenoids, α-carotene, lutein, β-carotene, and violaxanthin were the most abundant in both control and transgenic lines (Figure 3), while zeaxanthin and neoxanthin were below detection. Cell lines expressing *CrOR^WT^* or *CrOR^His^* accumulated significantly more of each detectable carotenoid than the control line. The α-carotene levels were up to 2.5- and 3.7-fold higher than those of the control in *CrOR^WT^* and *CrOR^His^* lines, respectively, with similar increases observed for lutein (2.3 and 2.8-fold higher), β-carotene (2.2 and 3.1-fold higher) and violaxanthin (2.3 and 3.4-fold higher) (Figure 3).

### 3.3. Both CrOR^WT^ and CrOR^His^ dramatically alters the expression levels of carotenogenic genes

We performed q-PCR to determine whether high-carotenoid accumulation in transgenic algae is correlated with higher expression of the genes related to the carotenoid biosynthesis pathway, including *DXS1, PSY1, LCY-β, LCY-ε, CYP97C3, ZEP*, and *NXS*. As shown in Figure 4, all key genes in carotenoid biosynthesis pathway except for *NXS*, were up-regulated by roughly two-fold in both *CrOR^WT^* and *CrOR^His^* lines relative to controls, with slightly higher gene expression in *CrOR^His^* lines. These results suggest that the observed higher levels of carotenoid accumulation in the transgenic lines can be attributed at least in part to up-regulated expression of carotenogenic genes caused by overexpression of *OR*.

### 3.4. CrOR overexpression alters C. reinhardtii cell size and chloroplast shape

In various higher plant species, *OR* mutant alleles have been shown to trigger plastid differentiation and accumulation of carotenoids (Lopez et al., 2008; Yuan et al., 2015a; Yazdani et al., 2019) and also increase the plastid size (Yuan et al., 2015a; Yazdani et al., 2019). We hypothesized that the *CrOR* overexpression may also change the plastid size in *C. reinhardtii*, therefore we used confocal microscopy to compare the size of the plastids within transgenic cells as well as measure overall cell size. We observed that cells of both *CrOR^WT^* and *CrOR^His^* lines were larger than the non-transgenic cells (Figure 5A). The average projected cell area of *CrOR^WT^* and *CrOR^His^* cells was approximately 45% and 73% larger, respectively, than that of the control cells (Figure 5B). Calculated estimates of cell volume, using the average radii from the projected area and an assumed spherical cell, reveal that *CrOR^WT^* and *CrOR^His^* cells were up to 1.7- and 2.3-fold larger on a volume basis, respectively, than the control cells (data not shown), which contributes to, but does not entirely explain, the elevated per cell carotenoid levels noted above.

Noticeably, the chloroplast size and shape in *CrOR^WT^* and *CrOR^His^* also showed some differences from the control line. While the chloroplasts in control cells exhibited a characteristic horse shoe shape, they were much larger in size and they lost the regular shape in both *CrOR^WT^* and *CrOR^His^* overexpressing cells. The increase in cell size and loss of characteristic chloroplast shape was more pronounced in *CrOR^His^* overexpressing lines (Figure 5A). These results clearly show that as in higher plants the OR protein can influence plastid size and can also increase algae cell size.

### 3.5. Transgenic C. reinhardtii cells showed enhanced tolerance to salt and oxidative stress

To investigate whether carotenoid-enhanced lines are better able to grow under abiotic stress conditions, *CrOR^WT^* and *CrOR^His^* transgenic lines were grown on TAP medium supplemented with various concentrations of either NaCl or paraquat and their growth rate was measured. Paraquat induces the production of superoxide radical, which has adverse effects on the electron transfer machinery in the mitochondrion and chloroplast and can therefore be used to test cellular resistance to oxidative stress (Bowler et al., 1991; Tunc-Ozdemir et al., 2009).

The growth rates of *CrOR^WT^* and *CrOR^His^* lines were approximately 35% and 72% higher than those of control cells in medium containing 50 and 100 mM NaCl which indicated that both *CrOR^WT^* and *CrOR^His^* lines were significantly more tolerant of salt stress than the control and the *CrOR^His^* line was significantly more tolerant than the *CrOR^WT^* line (Figure 6A). In the presence of 200 mM NaCl, culture fluorescence decreased over time in all genotypes; however, *CrOR^His^* line showed less fluorescence decline rate in comparison to *CrOR^WT^* and control lines (Figure 6B).

**Figure 6:**
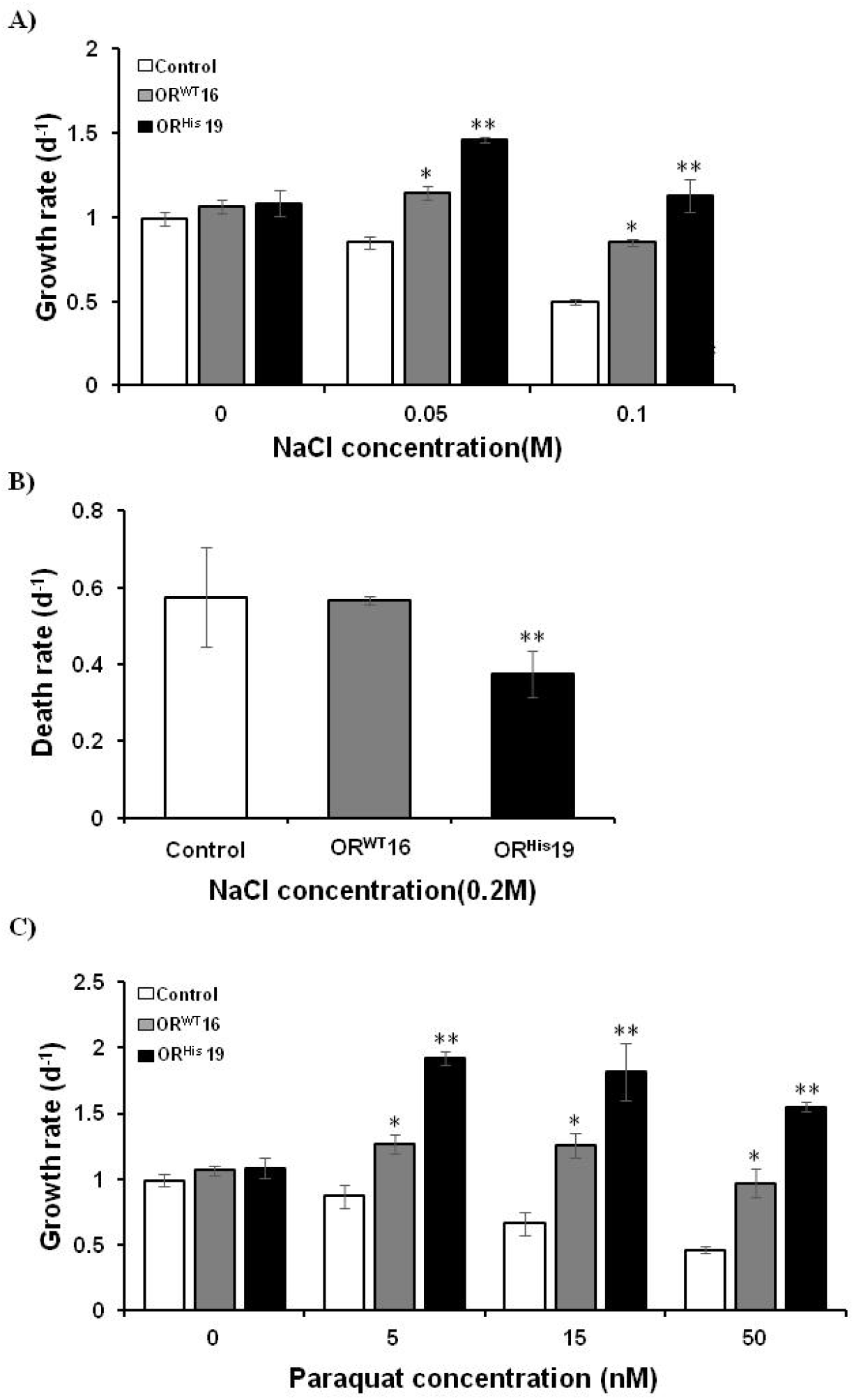
Effects of various concentrations of salt (A and B) and paraquat (C) on growth of *CrOR* transgenic lines. Bars denote the mean of three biological replicates ± SE. * indicates significant difference at *P*<0.05.

A similar result was observed in the paraquat experiments. The control cell line exhibited steadily lower growth rates with increasing paraquat concentration reaching 50% of its baseline rate with 50 nM added paraquat. In contrast, both *CrOR^WT^* and *CrOR^His^* lines maintained or exceeded baseline growth rates in response to added paraquat (Figure 6C). The end result is that growth rates for 5, 15, and 50 nM added paraquat for the *CrOR^WT^* and *CrOR^His^* lines were approximately 1.5-, 1.9-, and 2.1-fold higher and 2.2-, 2.8-, and 3.4-fold higher, respectively, than the control (Figure 6C). Similar to the salt stress experiment, the growth rate in *CrOR^His^* line was significantly higher than the *CrOR^WT^* line in all three concentrations of paraquat (Figure 6C). These results clearly indicate that carotenoid accumulation mediated by *OR* overexpression helps the *C. reinhardtii* to cope with abiotic stress conditions.

### 3.6. OR enhances the ABA accumulation in transgenic lines under oxidative stress condition

In higher plants, ABA is known as the stress hormone and plays an important role in the response to various stress conditions including biotic and abiotic stresses (Zhang, 2014). It has been demonstrated that algae are also able to synthesize ABA; however, its physiological functions have not been fully deciphered (Tietz and Kasprik, 1986; Tietz et al., 1989; Yoshida et al., 2003).

Because ABA is synthesized from the carotenoid cleavage pathway, the higher levels of carotenoids in the transgenic lines raised the question of whether they might also accumulate more ABA in comparison to the control line. Indeed, both *CrOR^WT^* and *CrOR^His^* lines accumulate higher levels of ABA than the control line (no added paraquat; Figure 7A). The amount of ABA normalized to algal DW was approximately 1.9- and 3.1-fold higher in *CrOR^WT^* and *CrOR^His^* lines, respectively. Consistent with these results, the expression level of the *NCED* gene was roughly double the control in *CrOR^WT^* and *CrOR^His^* lines (Figure 7B).

**Figure 7:**
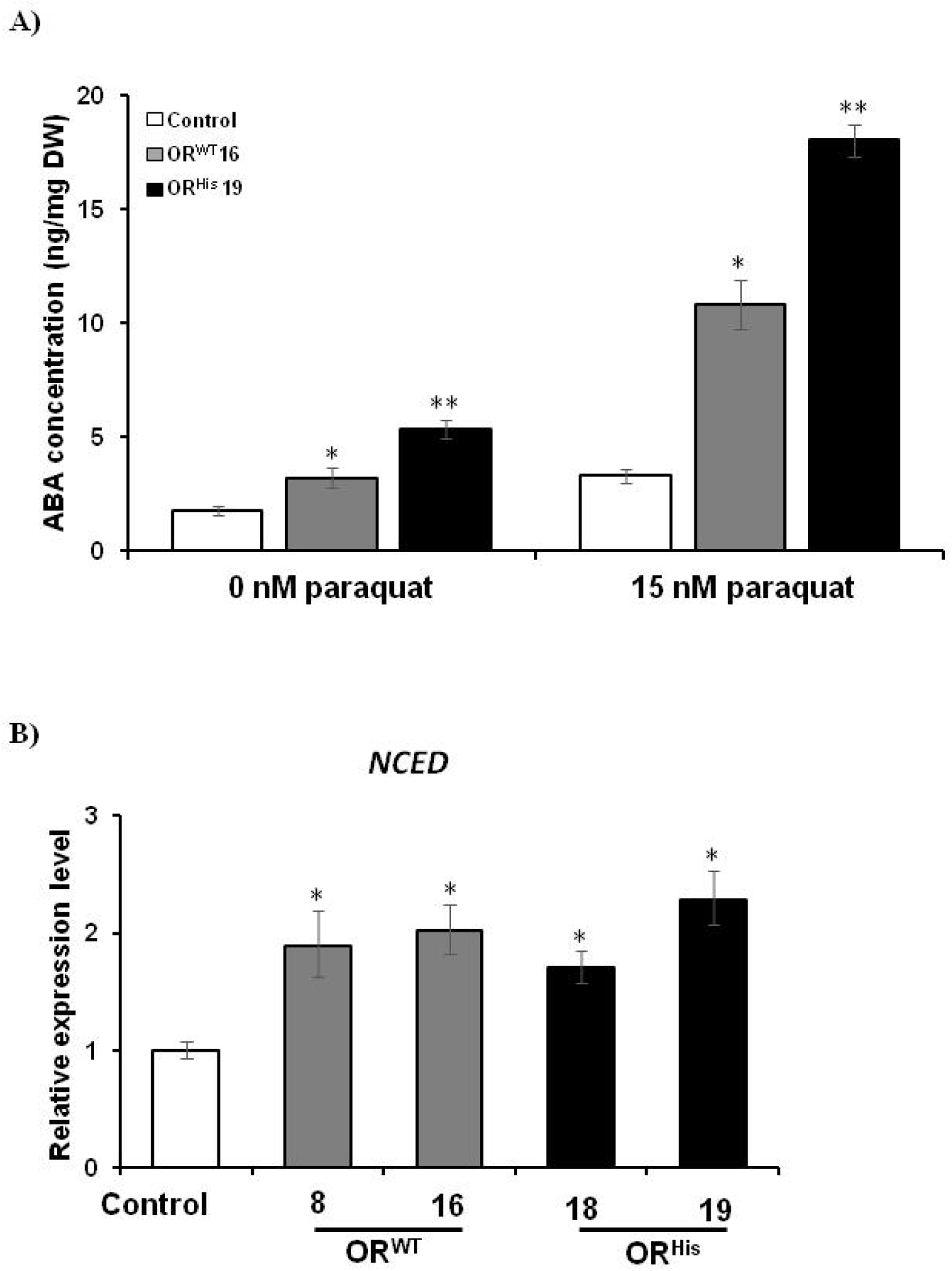
The effects of *CrOR* overexpression on ABA levels in transgenic cells. A, ABA concentration in transgenic lines under paraquat atress. B, expression levels of *NCED* gene in *OR* transgenes under normal growth condition. Bars denote mean of tree biological replicares ± SE. * indicates significant difference at *P*<0.05.

We also examined whether transgenic algae are capable of synthesizing more ABA under conditions of abiotic stress by measuring ABA in cells exposed to 15 nM paraquat. In response to the addition of paraquat, the high-carotenoid lines accumulated significantly more ABA than the control line, approximately 3.3- and 5.5-fold higher in *CrOR^WT^* and *CrOR^His^* lines, respectively (Figure 7A). Furthermore, in comparison to untreated cells (0 nM paraquat), the ABA levels in control and transgenic lines treated with 15 nM paraquat were about 2- and 3.5-fold higher (Figure 7A). These results clearly indicate that higher levels of endogenous ABA in algal cells is associated with a higher tolerance to abiotic stress conditions.

## 4. Discussion

In higher plants, overexpression of both wild type and mutated *ORANGE* gene has been used as a valuable genetic tool to elevate carotenoid levels in various species (Sun et al., 2018; Yazdani et al., 2019). For instance, in sweetpotato and *Arabidopsis* calli (Kim et al., 2013; Yuan et al., 2015a; Kim et al., 2019), rice and maize grains (Bai et al., 2016; Berman et al., 2017), and tomato fruit (Yazdani et al., 2019) overexpression of the wild type *OR* gene, in some cases with carotenogenic genes, has resulted in plant lines that accumulated 2- to 17-fold higher total carotenoids levels as compared to control lines. In many cases, plants overexpressing the *OR^His^* were found to synthesize and accumulate carotenoids at even higher levels compared to the plants overexpressing the *CrOR^WT^*. For instance, an approximately 13-fold increase in total carotenoid in sweetpotato calli (Kim et al., 2019), a 4-fold increase of total carotenoid content in *Arabidopsis* calli (Yuan et al., 2015a), and a 2-fold increase of total carotenoid at red stage of tomato fruit (Yazdani et al., 2019) has been reported in tissues overexpressing *OR^His^* compared to those overexpressing *OR^WT^*.

Consistent with these results, we observed a similar trend in total carotenoid levels in *C. reinhardtii* transgenic lines, in which total accumulated carotenoid was dramatically boosted in both *CrOR^WT^* and *CrOR^His^* overexpressing lines as compared to the control (Figure 3). While higher plants overexpressing the *OR* gene and especially the mutated *OR* preferentially accumulate lutein and β-carotene as the major forms of the carotenoid in their plastids (Lu et al., 2006; Kim et al., 2013; Yuan et al., 2015a; Yazdani et al., 2019), in our study, violaxanthin and α-carotene were also upregulated in response to both *OR* variants in *C. reinhardtii* transgenic lines (Figure 3). Enhanced accumulation of all four carotenoids could be the consequence of the upregulation of the MEP pathway genes including DXS as well as other relevant carotenogenic genes, specifially *PSY*, in both *CrOR^WT^* and *CrOR^His^* lines (Figure 4). In previous studies of higher plants, the expression of carotenoid biosynthetic genes is not significantly affected by *OR* expression (Lopez et al., 2008; Zhou et al., 2015; Yuan et al., 2015a; Kim et al., 2019). Our results suggest a new role for OR and show that unlike higher plants, OR in microalgae is able to transcriptionally regulate genes involved in several steps of carotenoid biosynthesis.

Carotenoids are essential for photosynthesis as they enable assembly of the photosynthetic machinery (Pogson et al., 2005). They also function as potent antioxidants quenching triplet chlorophyll, scavenging ROS to protect membranes and proteins, and reducing the detrimental effects of excess energy in the chloroplasts by non-photochemical quenching which is mediated by xanthophyll carotenoids (Pogson et al., 2005; Alboresi et al., 2011; Domonkos et al., 2013). In our study, *CrOR^WT^* and *CrOR^His^* transgenic lines displayed an improved tolerance to salt and oxidative stresses (Figure 6 A, B and C), perhaps in part mediated by the overaccumulation of carotenoids in these lines. These results are consistent with the results previously reported in various plant species in which *OR* transgenes, especially plants overexpressing the mutated *OR*, were more tolerant against various abiotic stress conditions such as salt (Kim et al., 2013; Kim et al., 2019), paraquat (Goo et al., 2015), drought (Wang et al., 2015; Cho et al., 2016), and heat (Park et al., 2016; Kim et al., 2019). Additionally, Park et al. (2016) demonstrated that under heat and oxidative stresses, sweetpotato OR protein (IbOr) directly interacts with PSY enzyme and stabilizes it by chaperone activity. These results suggest dual functions for the OR protein in protecting cells against abiotic stresses by regulating carotenoid biosynthesis and PSY protein stabilization.

Overexpression of the *OR^WT^* and *OR^His^* gene in higher plants results in larger plastids and the mutant *OR^His^* gene also triggers plastid differentiation into chromoplasts (Lopez et al., 2008; Yuan et al., 2015a; Yazdani et al., 2019). Consistent with these results, our study showed that overexpression of *CrOR^WT^* and *CrOR^His^* genes increased the plastid size, changed its shape and also increased cell size (Figure 5 A and B). Yuan et al. (2015a) used TEM to show that overexpression of *AtOR^WT^* resulted in larger plastoglobuli in *Arabidopsis* calli, whereas overexpression of the *AtOR^His^* gene resulted in the accumulation of large membrane stacks in plastids and a loss of plastoglobuli. The larger plastids observed in our *CrOR^WT^* and *CrOR^His^* transformants appear morphologically similar but lack of resolution in our images limits our ability to conclude if one of these same mechanisms led to the observed plastid enlargement and subsequently to the formation of larger cells.

ABA is phytohormone in higher plants that is synthesized in response to stress and it is known to positively regulate cell growth and elongation. For example, Ghassemian et al. (2000) showed that ABA at low concentrations promotes root growth in *Arabidopsis* and Fricke et al. (2004) demonstrated that ABA can resume leaf growth and cell elongation in barley under salinity stress. The role played by ABA in microalga is less well understood but ABA has been shown to have dramatic effects on cell morphology. In *H*. *pluvialis*, the application of ABA changed vegetative cells into red cyst cells (Kobayashi et al., 1997). In *C. reinhardtii*, exogenous ABA inhibited cell division which resulted in increase of cell size (Park et al., 2013) and in the unicellular red algae *Cyanidioschyzon merolae*, application of exogenous ABA arrested cell cycle and prevented algae cell proliferation (Kobayashi et al. 2016). Therefore, the larger cell size observed in our *CrOR^WT^* and *CrOR^His^* lines may alternatively be due to the over-production of ABA in these cells which inhibits cell division and ultimately the accumulation of larger cells.

Growth and development in higher plants are mainly regulated by internal and environmental cues, and ABA is an important regulator to coordinate growth and development in response to environmental signals (Xiong and Zhu, 2003). ABA is well-known as an important stress hormone, which helps plants to adapt and cope with various abiotic stresses such as salt, drought, low temperature, and osmotic stresses (Giraudat et al., 1994; Leung and Giraudat, 1998). In *C. reinhardtii*, the addition of ABA to the growth medium reduces the adverse effects of salt, osmotic, and paraquat stresses (Yoshida et al., 2003; Yoshida et al., 2004). Moreover, studies show that the application of exogenous ABA in algae has protective effects against ROS by enhancing the activity of antioxidant enzymes such as catalase, ascorbate peroxidase, and superoxide dismutase (Yoshida et al., 2003; Yoshida et al., 2004; Guajardo et al., 2016). These results indicate that ABA plays a key role in combating abiotic stresses not only in terrestrial plants, but also in microalgae.

Algae can convert carotenoids into ABA, which can function as a signaling molecule under oxidative stress (Yoshida, 2005; Kobayashi et al., 2016). In the present study, we showed that *CrOR* overexpressing lines accumulate higher levels of ABA than the control line when subjected to oxidative stress caused by paraquat (Figure 7) and these cells are more resistant to abiotic stress. To the best of our knowledge, this is the first report regarding the effect of increasing endogenous ABA on stress resistance in *C. reinhardtii*. The elevated levels of ABA in the *CrOR* transgenic lines may be a consequence of the overaccumulation of its precursor, i.e. carotenoids, in these cells, but in addition, we noted upregulation of the *NCED* gene. In *C. merolae*, a loss-of-function mutation in the *NCED* gene eliminated ABA biosynthesis even under salt stress, demonstrating that *NCED* is necessary for ABA biosynthesis (Yoshida et al., 2016). The evidence presented herein suggests a novel role for OR gene in triggering stress resistance in *C. reinhardtii*, and support the fact that similar to higher plants, ABA in *CrOR^WT^* and *CrOR^His^* transgenic cells may function as a signaling molecule to induce the activity of antioxidant enzymes under unfavorable conditions; however, the exact molecular mechanisms of these responses in algae require further investigations.

## Acknowledgement

This work was supported by discretionary funding to BAA from Cornell’s College of Agriculture and Life Science.

## Conflicts of interest

The authors declare that they have no competing interests.

## Authors’ contributions

MY and BAA conceived and designed the experiments. MY performed the experiments and wrote the manuscript. TLF and TWT helped and guided carotenoid analysis by UPC^2^. MGC aided with algae cell growth assay. MY and BAA analyzed the results. All authors contributed to the final manuscript.

---

**Supplemental Figure S1:**
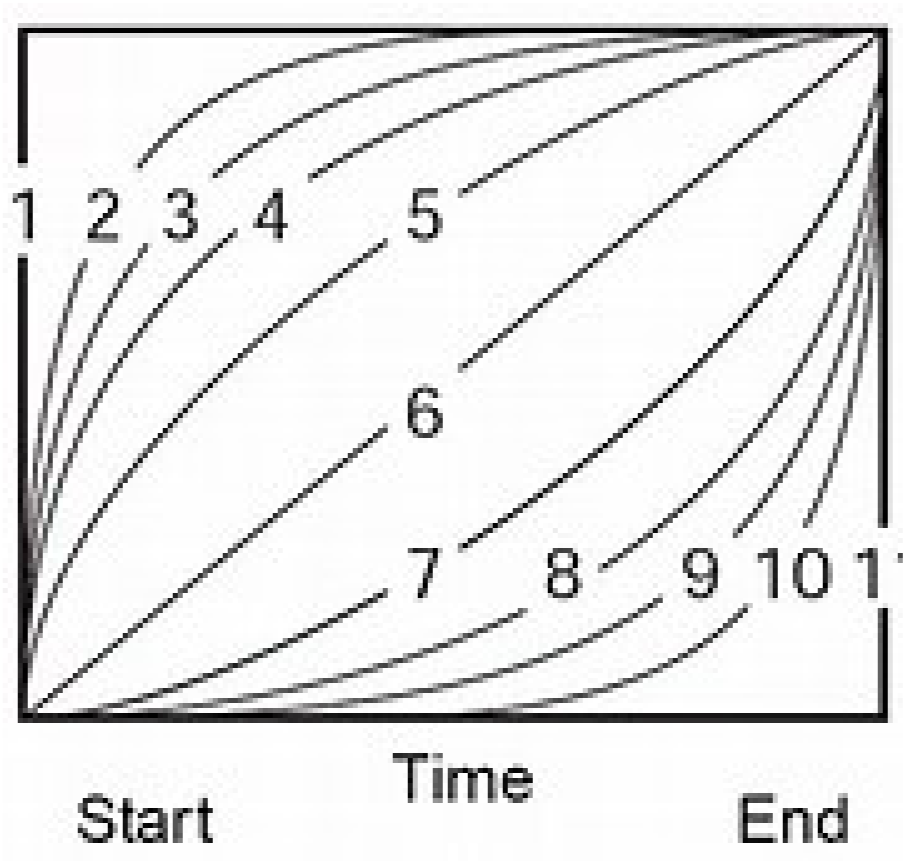
Waters UPC2 gradient curves

**Supplemental Table S1:**
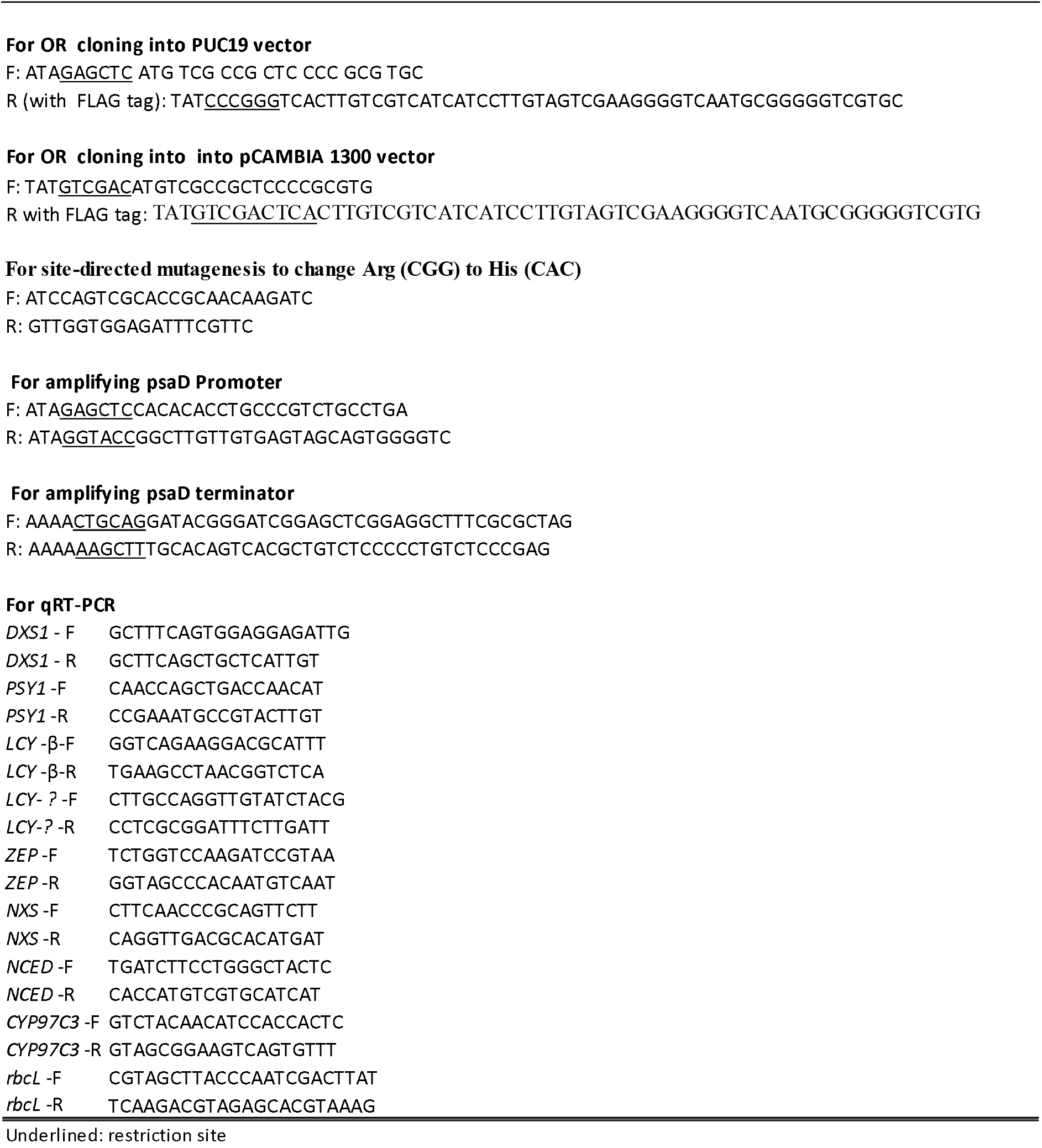
List of the primers used in this study

**Supplemental Table S2:** Waters UPC2 gradient conditions

|  | Time (min) | Flow (ml/min) | % A (Co2) | % B (Methanol) | curve |
| --- | --- | --- | --- | --- | --- |
| 1 | Initial | 1 | 99 | 1 | Initial |
| 2 | 7.5 | 1 | 80 | 20 | 10 |
| 3 | 10 | 1 | 80 | 20 | 6 |
| 4 | 12 | 1 | 80 | 20 | 6 |
| 5 | 15 | 1 | 99 | 0 | 1 |

**Supplemental Table S3:** ABPR pressure gradient setting

|  | Time (min) | Pressure (psi) |
| --- | --- | --- |
| 1 | Initial | 3750 |
| 2 | 7 | 3750 |
| 3 | 10 | 3000 |
| 4 | 12 | 3000 |
| 5 | 14 | 3750 |

**Supplemental Table S4:** Carotenoids optimal wavelengths and elution times by UPC2

| Compound | wavelength (nm) | Retention Time (min) |
| --- | --- | --- |
| α-carotene | 433 | 5.28 |
| β-carotene | 435 | 5.97 |
| Lutein | 431 | 11.28 |
| Violaxanthin 1 | 417 | 10.66 |
| Violaxanthin 2 | 417 | 11.1 |
| Violaxanthin 3 | 417 | 11.59 |
| DIM | 282 | 8.79 |

## Notes

https://www.chlamyorangeprotein.com

### Competing Interest Statement

The authors have declared no competing interest.

